# *PLCG2* as a Risk Factor for Alzheimer’s Disease

**DOI:** 10.1101/2020.05.19.104216

**Authors:** Andy P. Tsai, Chuanpeng Dong, Christoph Preuss, Miguel Moutinho, Peter Bor-Chian Lin, Nicole Hajicek, John Sondek, Stephanie J. Bissel, Adrian L. Oblak, Gregory W. Carter, Yunlong Liu, Gary E. Landreth, Bruce T. Lamb, Kwangsik Nho

## Abstract

Alzheimer’s disease (AD) is characterized by robust microgliosis and phenotypic changes that accompany disease pathogenesis. Indeed, genetic variants in microglial genes are linked to risk for AD. Phospholipase C *γ* 2 (PLCG2) participates in the transduction of signals emanating from immune cell-surface receptors that regulate the inflammatory response and is selectively expressed by microglia in the brain. A rare variant in *PLCG2* (P522R) was previously found to be protective against AD, indicating that *PLCG2* may play a role in AD pathophysiology. Here, we report that a rare missense variant in *PLCG2* confers increased AD risk (*p*=0.047; OR=1.164 [95% CI=1.002-1.351]). Additionally, we observed that *PLCG2* expression levels are increased in several brain regions of AD patients, correlating with brain amyloid deposition. This provides further evidence that *PLCG2* may play an important role in AD pathophysiology. Together, our findings indicate that *PLCG2* is a potential new therapeutic target for AD.

## Introduction

Late-onset Alzheimer’s disease (LOAD) is the most common form of AD, with symptoms typically arising after age 65 ^1^. Age, sex, and the apolipoprotein ε4 (*APOE* ε4) allele are the three greatest risk factors for LOAD ^2-4^. The risk for LOAD doubles every five years after age 65, and over 60% of AD patients are female ^2,5^. The mechanisms underlying the development of LOAD are incompletely understood, but recent large-scale genome-wide association studies (GWAS) have identified that more than 30 genetic variants are associated with LOAD ^6,7^.

Importantly, approximately 40% of the identified genes are immune- and microglia-related, suggesting that microglia are involved in modulating AD pathology ^8^. Among these microglia-related genetic factors, a rare variant in *PLCG2* (phospholipase C *γ* 2), P522R (rs72824905-G), was found to be associated with reduced LOAD risk ^9^. PLCG2 is a membrane-associated enzyme that catalyzes the conversion of phospholipid PIP2 (1-phosphatidyl-1D-myo-inositol 4,5-bisphosphate) to IP3 (1D-myo-inositol 1,4,5-trisphosphate) and DAG (diacylglycerol), which plays a crucial role in cell-surface receptor signal transduction ^10^. Specifically, *PLCG2* is necessary for immune cell function and is highly expressed in microglia. Genomic deletions (exon 19, Δ19, or exon 20 through 22, Δ20-22) or somatic mutations (R665W or S707Y) within the regulatory domain of *PLCG2* resulted in constitutive activation and resistance to treatment by inhibition of its upstream activator, BTK (Bruton’s tyrosine kinase), in leukemia ^11,12^. Interestingly, a missense M28L variant in *PLCG2* that does not alter the constitutive activity of the enzyme following Rac2 GTPase stimulation has been identified in BTK-inhibitor-resistant patients ^13^. BTK inhibition reduces phagocytosis and inhibits the uptake of synaptic structures in rodent microglia, indicating a role for the BTK-PLCG2 pathway in microglial function ^14^.

Here, we report that expression levels of *PLCG2* are increased in AD, and that the M28L variant is associated with elevated LOAD risk. Similarly, *Plcg2* is highly expressed in the brain of an AD mouse model. Importantly, upregulated expression of *PLCG2* is associated with increased amyloid plaques, which was validated in an amyloid mouse model. Elucidating the different effects of *PLCG2* variants on LOAD risk will expand our understanding of the function of microglia in AD pathophysiology.

## Results

### The *PLCG2* M28L Variant Confers a Higher Risk for LOAD

Using previously published large-scale GWAS summary statistics ^6^, we identified that the missense variant M28L in *PLCG2* is associated with LOAD. The minor allele (T) of M28L in *PLCG2* significantly increased risk for LOAD (p=0.047, OR=1.164 [95% CI=1.002-1.351]) (**Table 1**). PLCG2 possesses a set of core domains common to most other isoforms of PLC, including an N-terminal PH domain, two pairs of EF hands, a catalytic TIM barrel composed of the X- and Y-boxes, and a C2 domain (**Fig. 1a)**. The PLCG isozymes elaborate on this core with a unique array of regulatory domains, including a split PH (sPH) domain, two SH2 domains, and an SH3 domain. Two substitutions that modulate risk for LOAD, M28L, and P522R, were mapped onto a protein structure homology model of PLCG2 ^15^ (**Fig. 1b**). M28 is buried in the core of the PH domain but is not visible in a space-filling model (**Fig. 1c**). The PH domain is essential for membrane association of PLCG2.

**Table 1:** *PLCG2* M28L confers a higher risk of LOAD.

|  |
| --- |
| *PLCG2* M28L (rs61749044) |
| p=0.047 |
| OR=1.164 [95% CI=1.002-1.351] |

OR *odds ratio*, CI *confidence interval*

**Table 2:** Demographic information of the participants included in this study.

| Brain region | TCX |  | PHG |  |  | STG |  |  | IFG |  |  | FP |  |  | CER |  | DLPFC |  |
| --- | --- | --- | --- | --- | --- | --- | --- | --- | --- | --- | --- | --- | --- | --- | --- | --- | --- | --- |
| Diagnosis | CN | AD | CN | MCI | AD | CN | MCI | AD | CN | MCI | AD | CN | MCI | AD | CN | AD | CN | AD |
| No. of participants | 71 | 80 | 16 | 14 | 78 | 21 | 18 | 98 | 18 | 16 | 102 | 22 | 20 | 111 | 72 | 79 | 86 | 155 |
| Sex (Female/Male) | 35/36 | 49/31 | 11/5 | 7/7 | 54/25 | 16/5 | 11/7 | 65/33 | 12/6 | 7/9 | 69/33 | 16/6 | 11/9 | 75/36 | 35/37 | 47/32 | 47/39 | 109/46 |
| Mean age at death (SD), years | 82.7 (8.5) | 82.6 (7.7) | 83.5 (8.9) | 80.6 (10.8) | 85.5 (6.1) | 83.8 (8.1) | 82.1 (10.4) | 84.5 (6.7) | 83.2 (8.6) | 80.8 (10.5) | 85.4 (6.0) | 83.1 (7.7) | 83.2 (9.8) | 85.4 (5.9) | 82.3 (8.3) | 82.5(7.7) | 83.4 (5.9) | 88.2 (3.1) |
| Mean RIN (SD) | 7.7 (1.0) | 8.6 (0.6) | 7.1 (0.9) | 6.7 (0.7) | 6.5 (0.9) | 6.4 (1.0) | 6.4 (0.8) | 6.3 (0.9) | 8.7 (1.4) | 8.8 (1.6) | 8.0 (1.8) | 6.8 (0.9) | 7.0 (1.0) | 6.8 (0.9) | 7.7 (1.0) | 8.4 (0.7) | 7.3 (1.0) | 6.9 (0.9) |
| APOE genotype (ε4+/-ε4-) | 62/9 | 38/42 | 14/2 | 9/5 | 57/21 | 17/4 | 12/6 | 65/33 | 16/2 | 8/8 | 72/30 | 19/3 | 13/7 | 70/41 | 62/10 | 38/41 | 77/9 | 91/64 |

TCX temporal cortex, PHG *parahippocampal gyrus*, STG *superior temporal gyrus*, IFG *inferior temporal gyrus*, FP *frontal pole*, CER *cerebellum*, DLPFC *dorsolateral prefrontal cortex*, CN *cognitively normal* , AD *Alzheimer’s disease*, MCI *mild cognitive impairment*, RIN *RNA integrity number*, APOE ε4+/- carriers and non-carriers of the APOE ε4 allele

**Fig 1.**
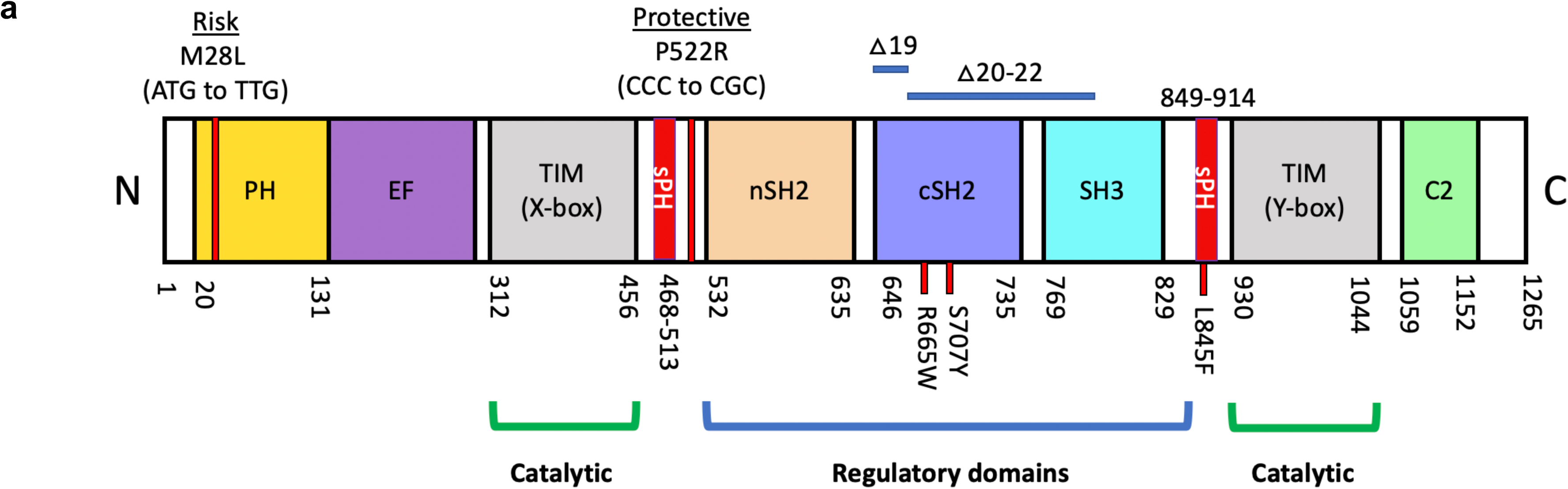
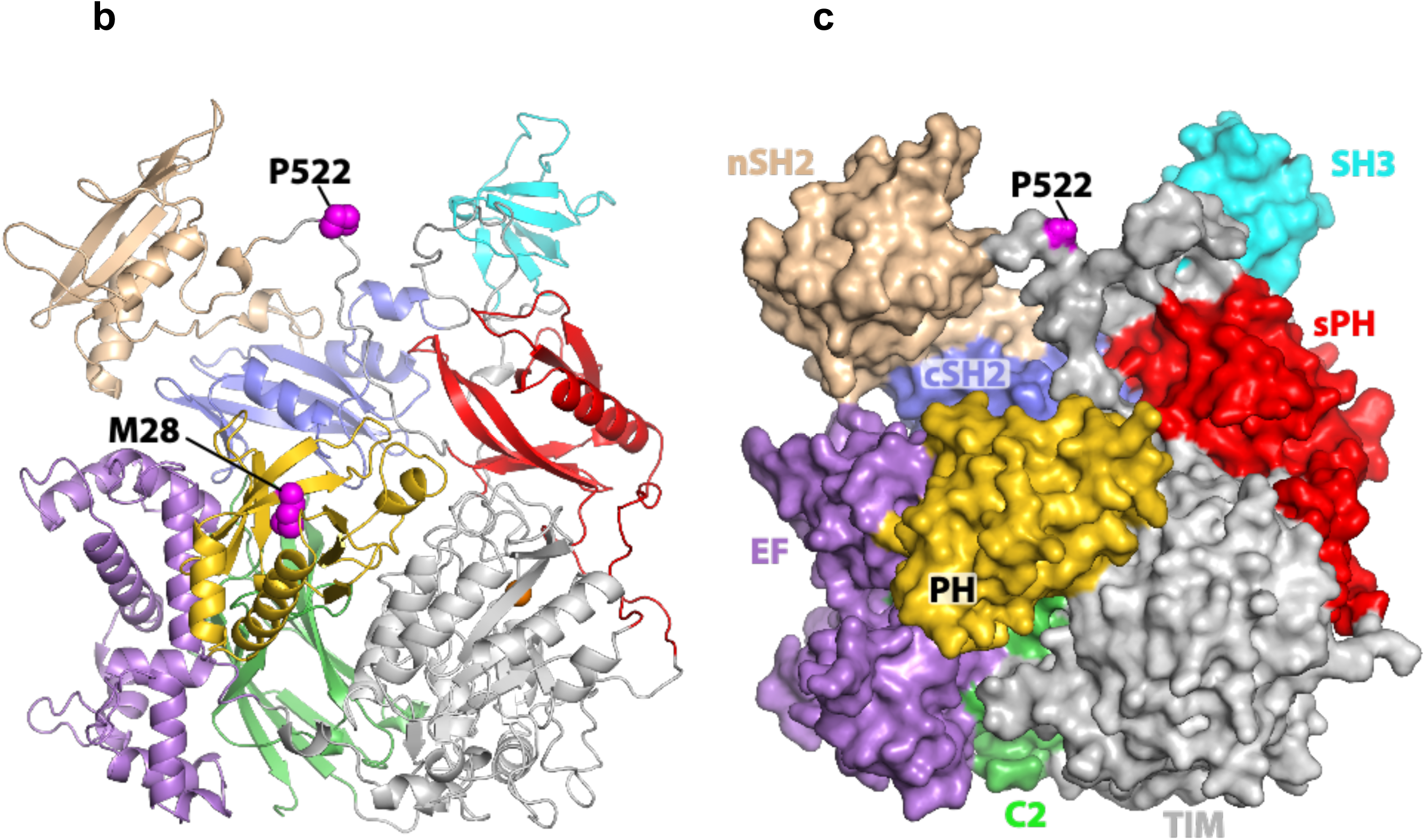
*PLCG2* M28L variant in PH domain. Domain architecture of *PLCG2* drawn to scale. Genomic deletions (exon 19, Δ19, or exon 20-22, Δ20-22) or somatic mutations (R665W or S707Y) in *PLCG2* within the regulatory domains are shown in the domain architecture. *PLCG2* M28L (risk) and P522R (protective) variants are shown in the domain architecture (a) and mapped onto the structure of PLCG2 (magenta spheres) in both homology model (b) and space-filling model (c). N *amino-terminus*, C *carboxyl-terminus*, PH *pleckstrin homology domain*, EF *EF hand motif*, TIM *TIM barrel*, sPH *split PH domain*, nSH2 *n- terminus Src Homology 2 domain*, cSH2 *c-terminus Src Homology 2 domain*, SH3 *SRC Homology 3 domain*, C2 *C2 domain*

### *PLCG2* Expression Levels are Increased in LOAD

We performed gene expression analysis using RNA-Seq data generated from seven brain regions. The demographic information of the participants included in this study is summarized in **Table 2**. We found that *PLCG2* was over-expressed in LOAD in the temporal cortex (logFC=0.27, p=4.56E-02; **Fig. 2a**), parahippocampal gyrus (logFC=0.55, p=1.74E-03; **Fig. 2b**), superior temporal gyrus (logFC=0.46, p=2.55E-02; **Fig. 2c**), and inferior prefrontal gyrus (logFC=0.36, p=1.38E-02; **Fig 2d**) with age and sex as covariates (**Table 3**). However, we did not find any diagnosis group differences in the cerebellum (**Fig. 2e**), frontal pole (**Fig. 2f**), and dorsolateral prefrontal cortex (**Fig. 2g**) (**Table 3**). With age, sex, and *APOE* ε4 carrier status as covariates, *PLCG2* remained over-expressed in LOAD in the parahippocampal gyrus (logFC=0.57, p=2.02E-03), superior temporal gyrus (logFC=0.42, p=4.95E-02), and inferior prefrontal cortex (logFC=0.33, p=2.60E-02) (**Table 3**). In the parahippocampal gyrus, superior temporal gyrus, and inferior prefrontal gyrus, *PLCG2* was over-expressed in LOAD with and without *APOE* ε4 carrier status as an additional covariate. We investigated whether *PLCG2* expression levels were associated with expression levels of microglia-specific marker genes (*AIF1* and *TMEM119*). Our analysis revealed that expression levels of *AIF1* and *TMEM119* were significantly associated with *PLCG2* expression levels in the frontal pole (*AIF1*: β=0.258, p=1.09E-06; *TMEM119*: β=0.518, p=1.28E-15), superior temporal gyrus (*AIF1*: β=0.33, p=8.98E-07; *TMEM119*: β=0.71, p<2E-16), parahippocampal gyrus (*AIF1*: β=0.419, p=5.43E- 07; *TMEM119*: β=0.806, p<2E-16), and inferior prefrontal gyrus (*AIF1*: β=0.257, p=6.15E-06; *TMEM119*: β=0.582, p=6.64E-15) (**Table 4**).

**Table 3.** *PLCG2* expression levels were increased in LOAD. shows the p-value for the gene expression analysis performed by limma using RNA-Seq data from AMP-AD Consortium.

| Brain Regions | TCX | PHG |  |  | STG |  |  | IFG |  |  |
| --- | --- | --- | --- | --- | --- | --- | --- | --- | --- | --- |
| Covariate | Age and Sex | Age and Sex |  |  | Age and Sex |  |  | Age and Sex |  |  |
| Contrast | CN vs. AD | CN vs. MCI | MCI vs. AD | CN vs. AD | CN vs. MCI | MCI vs. AD | CN vs. AD | CN vs. MCI | MCI vs. AD | CN vs. AD |
| logFC | 0.27056314 | 0.07196964 | 0.473066378 | 0.550085612 | 0.127793252 | 0.34416553 | 0.45549039 | 0.076802391 | 0.29211474 | 0.35622471 |
| p-value | 4.56E-02 | 9.99E-01 | 6.74E-02 | 1.74E-03 | 1.00E+00 | 1.29E-01 | 2.55E-02 | 1.00E+00 | 5.72E-02 | 1.38E-02 |
| Covariate | Age, Sex, and APOE ε4 status | Age, Sex, and APOE ε4 status |  |  | Age, Sex, and APOE ε4 status |  |  | Age, Sex, and APOE ε4 status |  |  |
| Contrast | CN vs. AD | CN vs. MCI | MCI vs. AD | CN vs. AD | CN vs. MCI | MCI vs. AD | CN vs. AD | CN vs. MCI | MCI vs. AD | CN vs. AD |
| logFC | 0.248525854 | 0.02041513 | 0.492722546 | 0.569868694 | 0.097484106 | 0.329619141 | 0.415487533 | 0.007469606 | 0.300240428 | 0.33069001 |
| p-value | 1.03E-01 | 1.00E+00 | 8.71E-02 | 2.20E-03 | 1.00E+00 | 1.73E-01 | 4.95E-02 | 1.00E+00 | 5.96E-02 | 2.60E-02 |

| Brain Regions | CER | FP |  |  | DLPFC |
| --- | --- | --- | --- | --- | --- |
| Covariate | Age and Sex | Age and Sex |  |  | Age and Sex |
| Contrast | CN vs. AD | CN vs. MCI | MCI vs. AD | CN vs. AD | CN vs. AD |
| logFC | 0.207937827 | -0.188185731 | 0.178098774 | 0.02365209 | -0.080816649 |
| p-value | 9.34E-02 | 9.85E-01 | 4.56E-01 | 9.67E-01 | 4.53E-01 |
| Covariate | Age, Sex, and APOE ε4 status | Age, Sex, and APOE ε4 status |  |  | Age, Sex, and APOE ε4 status |
| Contrast | CN vs. AD | CN vs. MCI | MCI vs. AD | CN vs. AD | CN vs. AD |
| logFC | 0.2365597 | -0.228761625 | 0.18490708 | 0.01861041 | -0.094639191 |
| p-value | 8.46E-02 | 9.53E-01 | 4.73E-01 | 9.82E-01 | 4.53E-01 |

*TCX* temporal cortex, *PHG* parahippocampal gyrus, *STG* superior temporal gyrus, *IFG* inferior temporal gyrus, *FP* frontal pole, *CER* cerebellum, *DLPFC* dorsolateral prefrontal cortex, *CN* cognitively normal , *AD* Alzheimer’s disease, *MCI* mild cognitive impairment, *logFC* log fold-change

**Table 4.** *PLCG2* expression levels were associated with amyloid plaque density and expression levels of microglia specific markers. shows the β coefficient (β), standard error (SE), and *p*-value for the association analysis between *PLCG2* expression levels and amyloid plaque density or expression levels of microglia specific markers, *AIF1* and *TMEM119* by general linear models.

| Brain Regions (MSBB) | Frontal Pole |  |  | Parahippocampal Gyrus |  |  | Superior Temporal Gyrus |  |  | Inferior Frontal Gyrus |  |  |
| --- | --- | --- | --- | --- | --- | --- | --- | --- | --- | --- | --- | --- |
| | $\beta$ | SE | *p*-value | $\beta$ | SE | *p*-value | $\beta$ | SE | *p*-value | $\beta$ | SE | *p*-value |
| Plaque Mean Density | 0.01 | 0.005 | 8.45E-02 | 0.027 | 0.006 | 5.22E-05 | 0.026 | 0.006 | 2.02E-04 | 0.017 | 0.006 | 2.73E-03 |
| *AIF1* | 0.258 | 0.051 | 1.09E-06 | 0.419 | 0.078 | 5.43E-07 | 0.33 | 0.063 | 8.98E-07 | 0.257 | 0.054 | 6.15E-06 |
| *TMEM119* | 0.518 | 0.057 | 1.28E-15 | 0.806 | 0.066 | <2E-16 | 0.71 | 0.064 | <2E-16 | 0.582 | 0.065 | 6.64E-15 |

**Fig 2.**
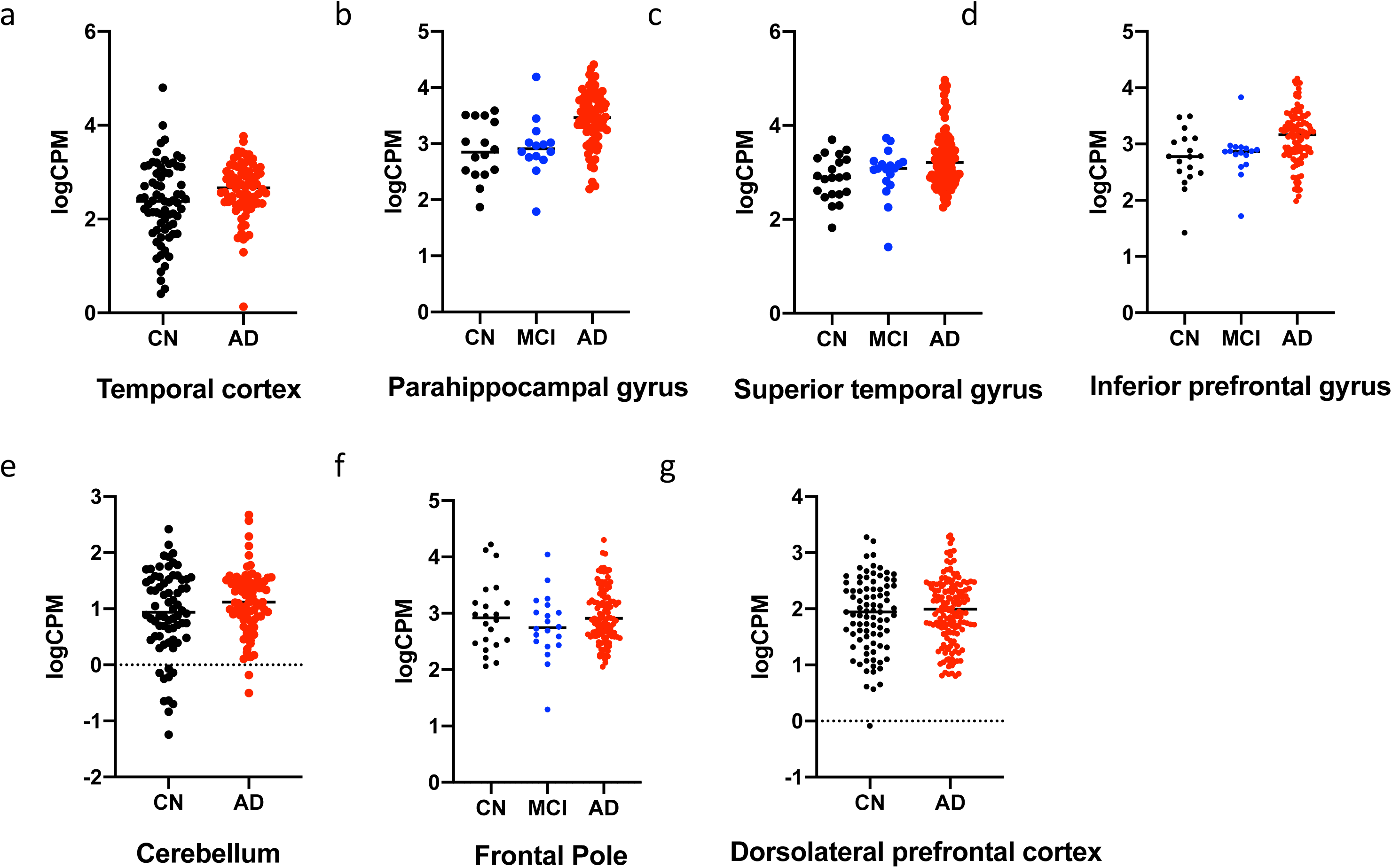
Relative quantification of *PLCG2* expression in the studied participants. Gene expression is showed as logCPM values. (a) Temporal cortex (TCX)-Mayo, (b) Parahippocampal gyrus (PHG)-MSBB, (c) Superior temporal gyrus (STG)-MSBB, (d) Inferior prefrontal gyrus (IFG)- MSBB, (e) Cerebellum (CER)-Mayo, (f) Frontal pole (FP)-MSBB, (g) Dorsolateral prefrontal cortex (DLPFC)-ROSMAP CN cognitively normal, AD Alzheimer’s disease, MCI mild cognitive impairment

### Increased *PLCG2* Expression Levels are Associated with Amyloid Plaque Density in the Human Brain

We performed an association analysis of *PLCG2* expression levels with mean amyloid plaque densities measured in four brain regions. Expression levels of *PLCG2* in the human brain was associated with amyloid plaques in three brain regions (**Table 4**). Increased *PLCG2* expression was associated with increased amyloid plaques in the parahippocampal gyrus (β=0.027, p=5.22E-05; **Fig. 3a**), superior temporal gyrus (β=0.026, p=2.02E-04; **Fig. 3b**), and inferior prefrontal cortex (β=0.017, p=2.73E-03; **Fig. 3c**), but not in the frontal pole (β=0.01, p=0.08; **Fig. 3d**).

**Fig 3.**
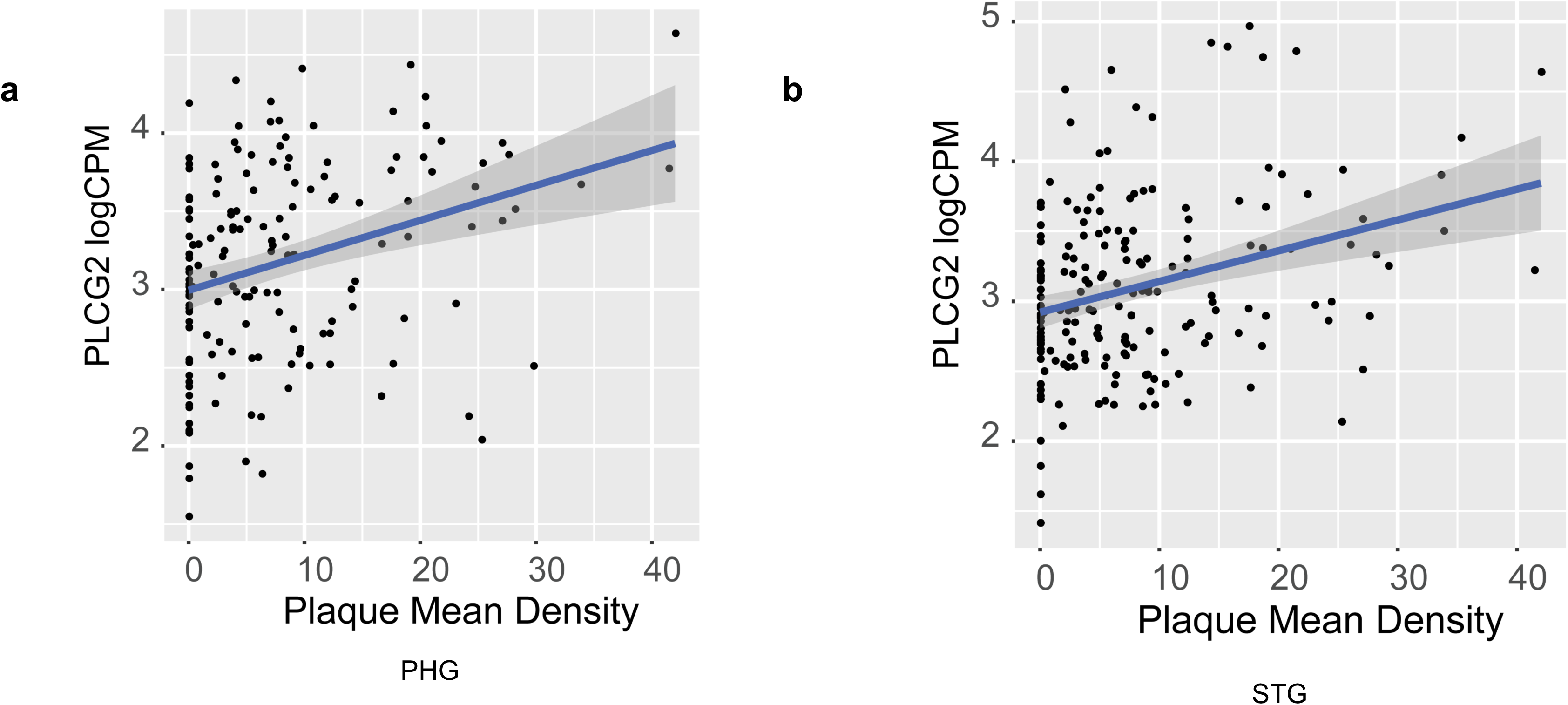
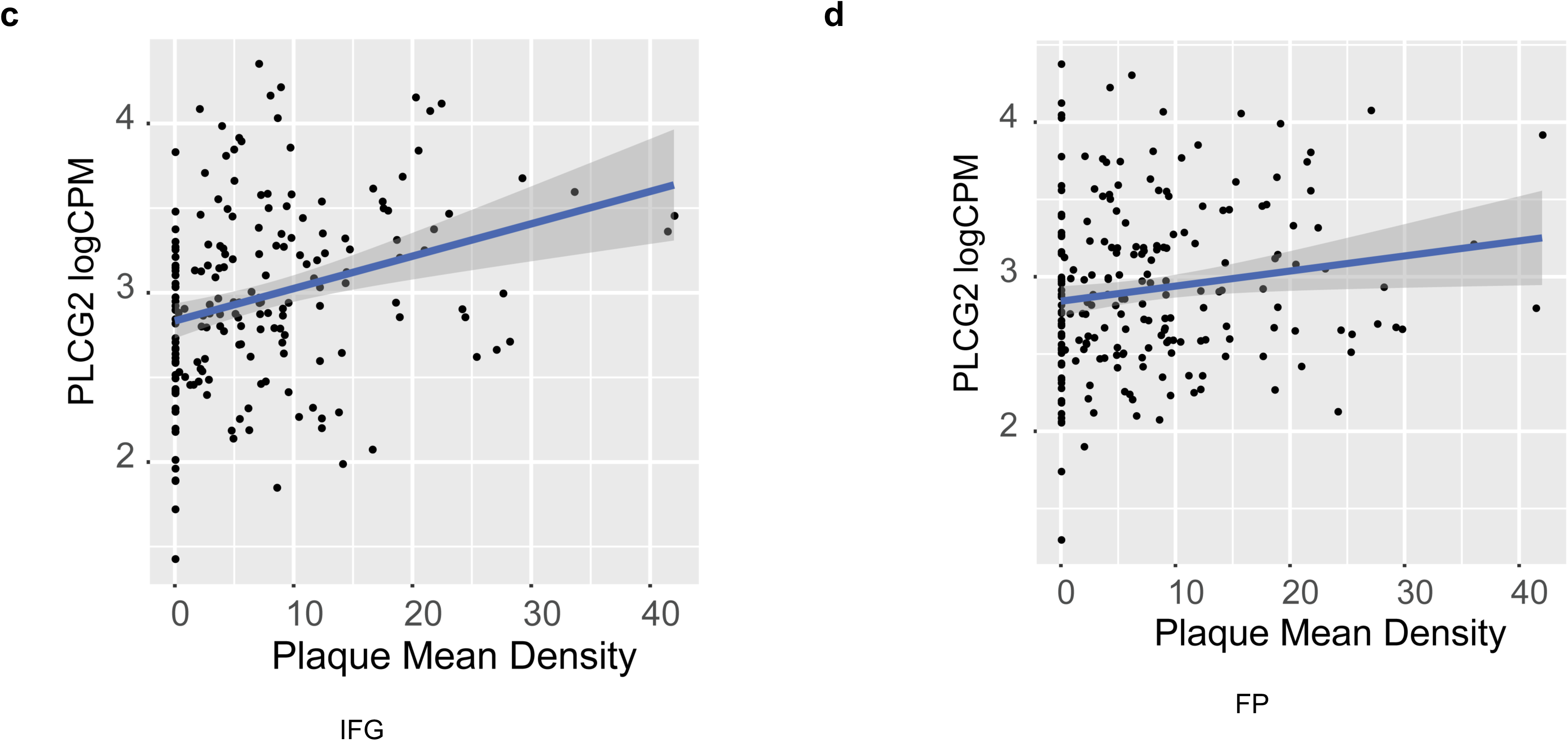
Associations of *PLCG2* expression with amyloid plaque mean density. The scatter plots show the positive association between *PLCG2* expression and plaque mean density in (a) parahippocampal gyrus, (b) superior temporal gyrus and (c) inferior prefrontal gyrus, and (d) frontal pole from MSBB cohort.

### *PLCG2* Expression Levels are Increased in a Mouse Model of Amyloid Pathology

*Plcg2* expression was increased in a mouse model of amyloid pathology, which is in accordance with the results from our analysis of human LOAD. In the 5xFAD mice, *Plcg2* expression was increased in the cortex (**Fig. 4a**) and hippocampus (**Fig. 4b)** of four-, six -and eight-month-old mice (4-month: 1.39-fold in the cortex, 1.37-fold in the hippocampus; 6-month: 2.37-fold in the cortex, 1.96-fold in the hippocampus; and 8-month: 2.43-fold in the cortex and 2.67-fold in the hippocampus) (**Fig. 4**). Furthermore, our analysis showed a disease progression-dependent increase in *Plcg2* expression in mice with amyloid pathology (**Fig. 4**).

**Fig 4.**
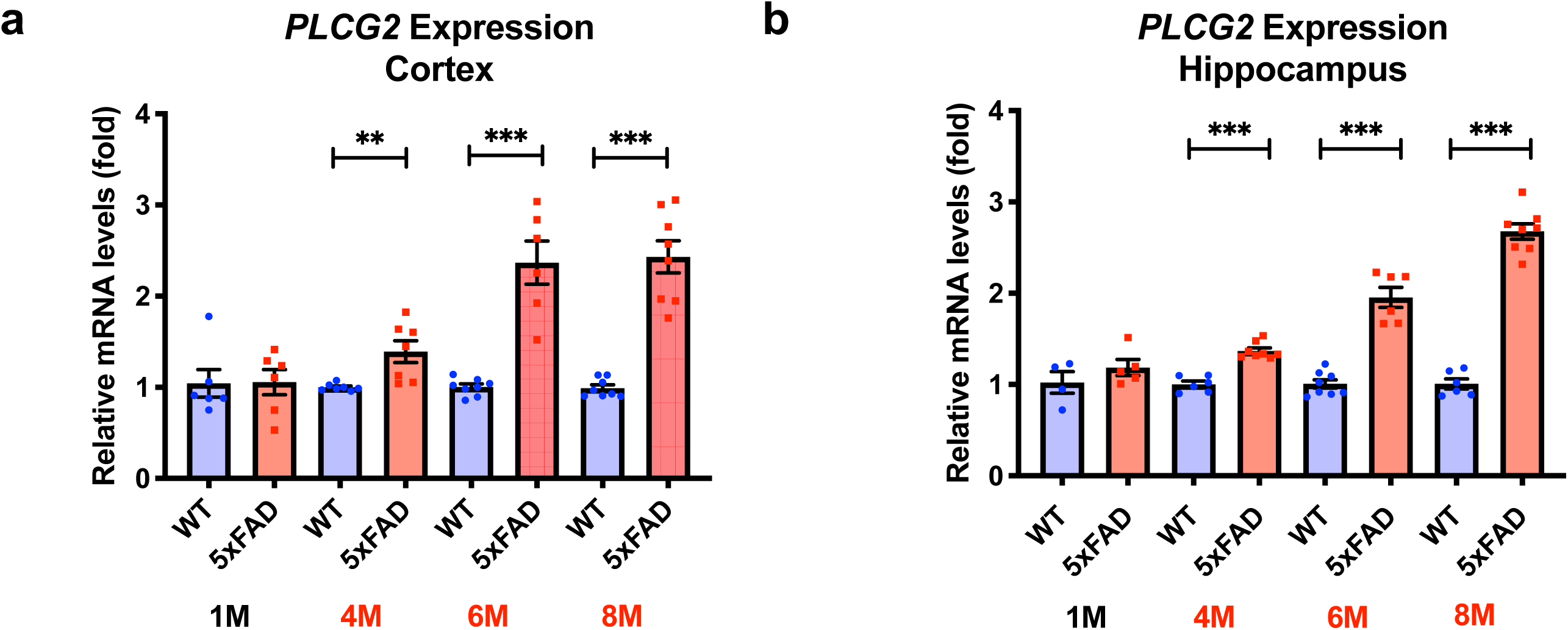
*Plcg2* expression levels were increased in a mouse model of amyloid pathology. *Plcg2* RNA levels were assessed in cortical and hippocampal lysates from 5xFAD mice. There were significant changes in the *PLCG2* gene expression in both cortex (a) and hippocampus (b) at 4 months, 6 months, and 8 months of age (n=4-8 mice). **p<0.01; ***p<0.001

### Common Single Nucleotide Polymorphisms in *PLCG2* are Associated with *PLCG2* Expression Levels

We performed an expression quantitative trait loci analysis (eQTL) analysis of *PLCG2* expression levels using common SNPs, including ±20kb from the gene boundary; minor allele frequency (MAF) > 5%, from whole-genome sequencing. The eQTL analysis identified several variants that are associated with *PLCG2* expression levels. In particular, rs4420523 was most significantly associated with *PLCG2* expression levels in the superior temporal gyrus (p=2.43E- 06, **Fig. 5a**). Individuals with minor alleles of rs4420523 (C) have higher *PLCG2* expression levels compared to those without minor alleles (**Fig. 5b**). The eQTL association of rs4420523 was replicated in the temporal cortex of the MAYO cohort (**Fig. 5c and Fig. 5d**). Furthermore, using the GTEx eQTL database, we found that rs4420523 is significantly associated with *PLCG2* expression levels in several brain regions and other organs (**Fig. 5e**).

**Fig 5.**
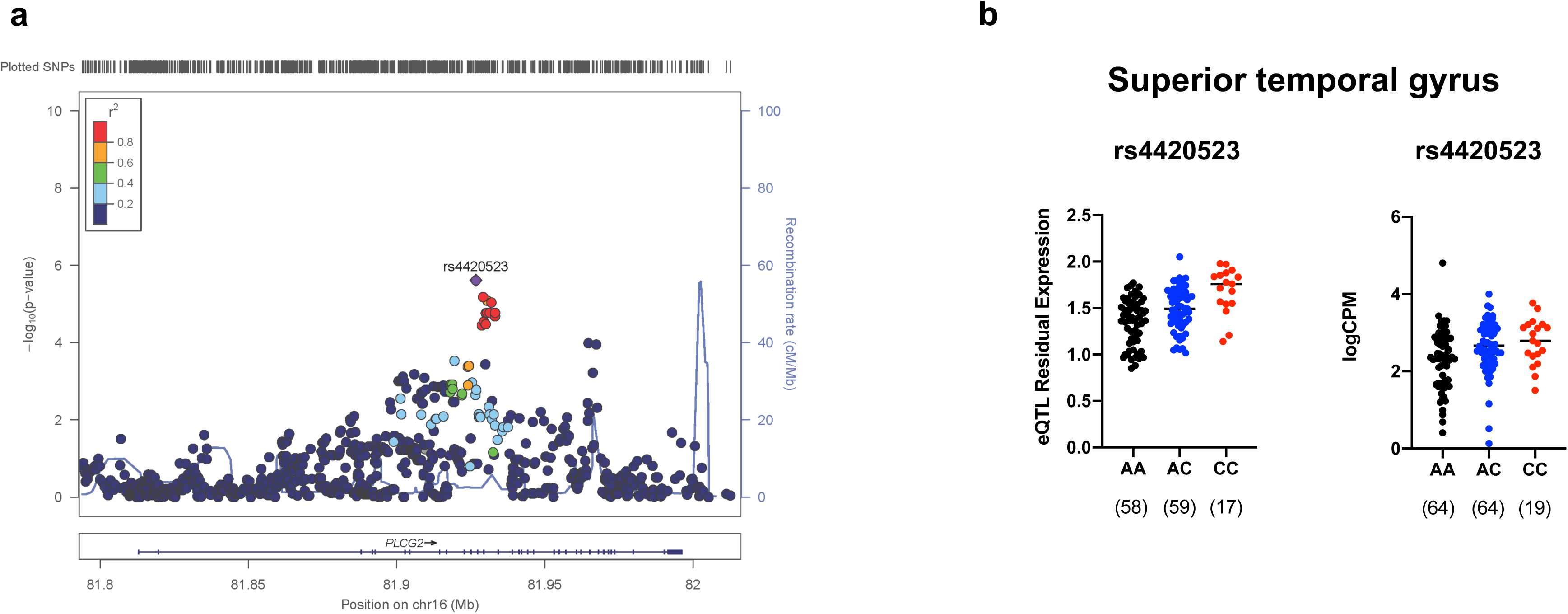
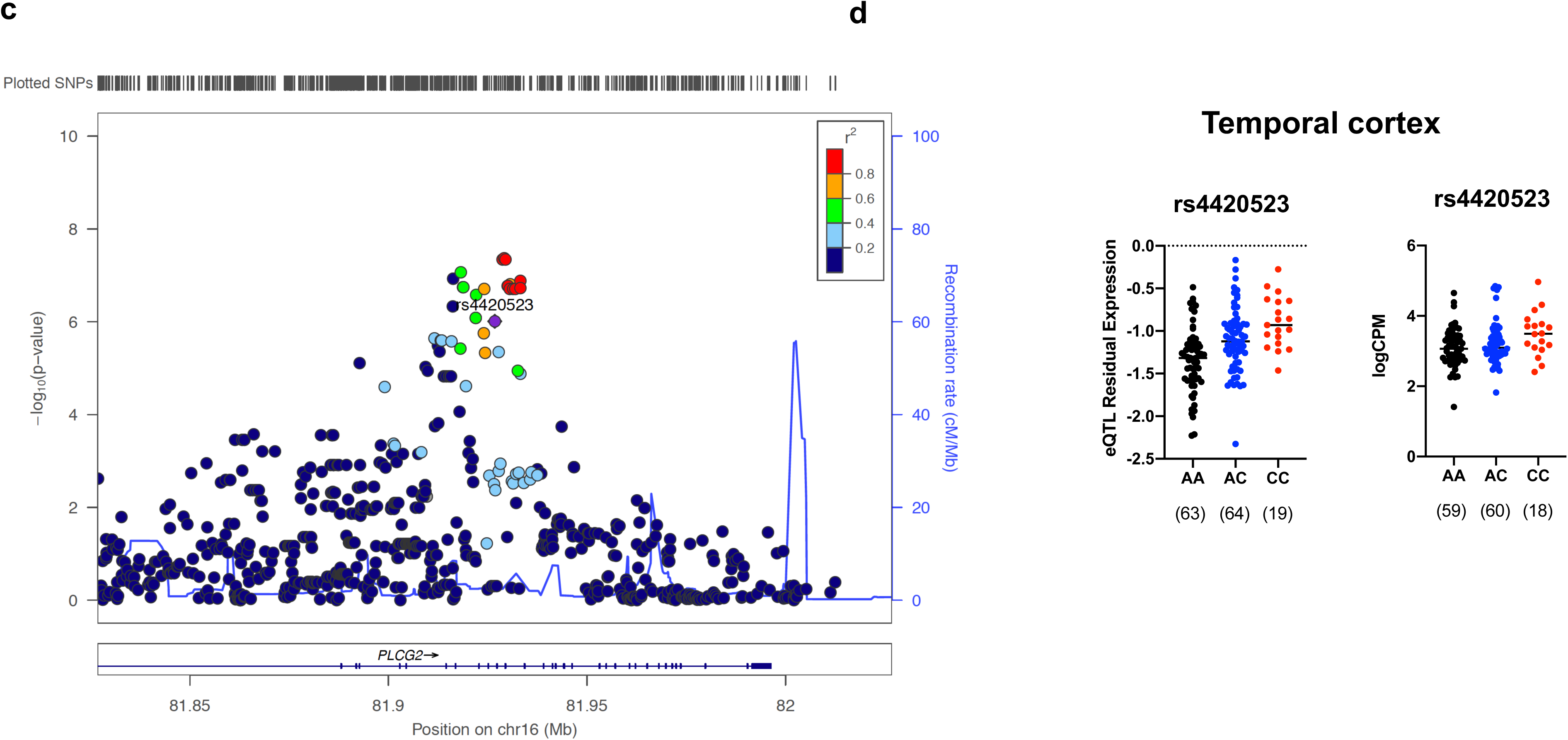
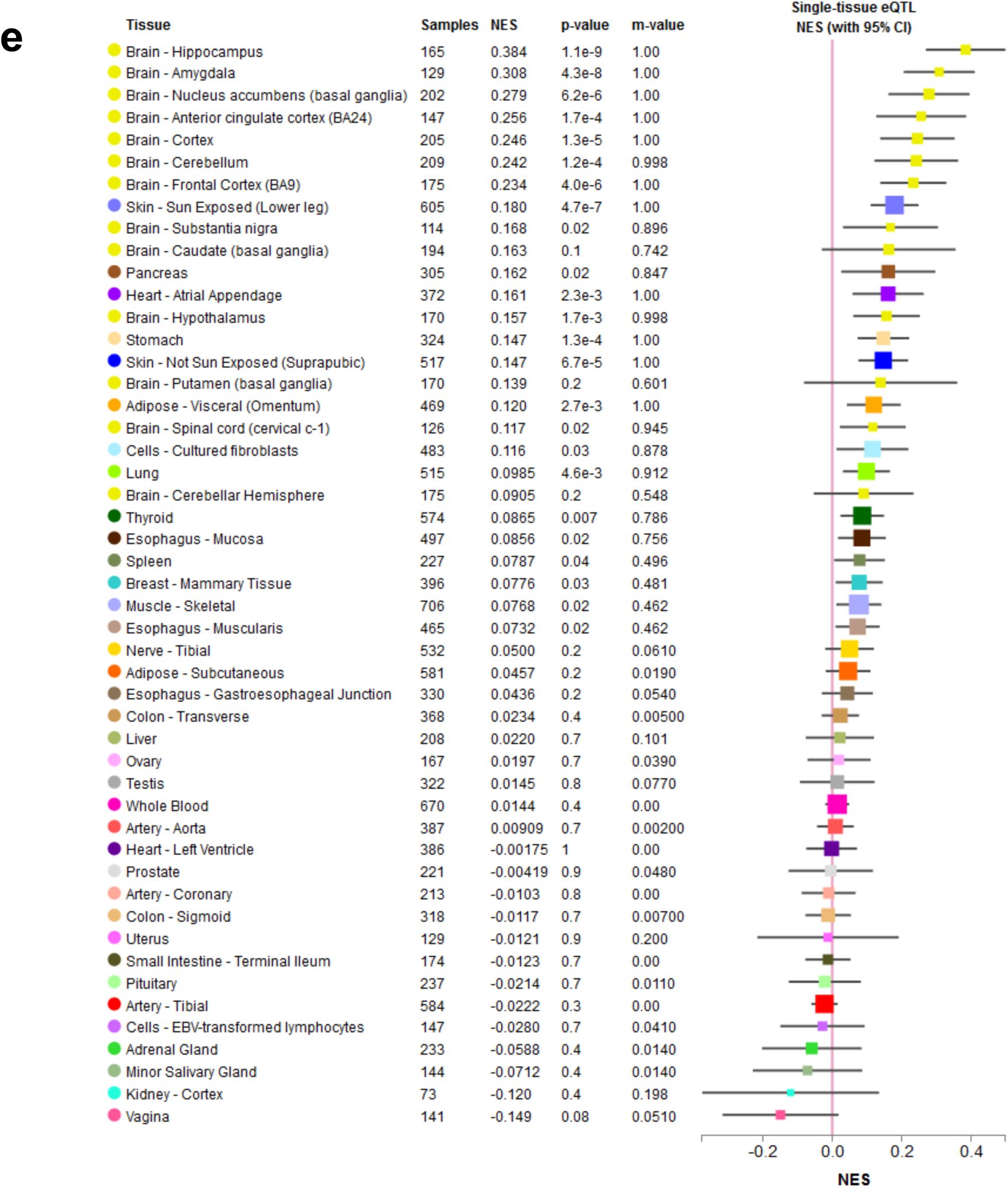
eQTL analysis of *PLCG2*. SNP positions on chromosome 16 and their color-coded association with *PLCG2* expression from (a) superior temporal gyrus and (c) temporal cortex are plotted by LocusZoom. The SNP (rs4420523) with lowest *p-*value is indicated. The *PLCG2* expression levels of participants with or without rs4420523 are showed as both eQTL residual expression and logCPM value from superior temporal gyrus (b) and temporal cortex (d). The numbers of participants are shown below the genotypes. The results of single-tissue eQTL analysis for rs4420523 are presented in (e). NES *normalized effect size*, chr16 *chromosome 16*, eQTL *expression quantitative trait loci analysis*, Major allele *A*, Minor allele *C*

## Discussion

In this study, we report the first evidence that a missense variant (M28L) in *PLCG2* is associated with increased risk for LOAD. The results indicated that different genetic variants in *PLCG2* may be either advantageous or deleterious in LOAD, suggesting a potentially important role of this gene in modifying AD risk. The M28L variant was previously found to be associated with leukemia and mechanistically linked to disease pathogenesis by its resistance to inhibition of its upstream kinase activator BTK ^13^. PLCG2 activity can be regulated by several mechanisms, including phosphorylation by kinases such as BTK^16,17^, PH domain-dependent regulation ^18-20^, SH2 domain-containing components regulation ^21^, and autoregulation of the intrinsic inhibition region ^21,22^. The methionine 28 is buried within the N-terminal PH domain, and the substitution likely leads to a less well-packed PH domain with a concomitant loss of domain stability. The instability of the PH domain may affect the protein-protein interactions that regulate enzymatic activity. Additionally, the conformational change may disrupt PLCG2-BTK or other protein interactions, leading to ibrutinib resistance. *PLCG2* is selectively expressed by microglia in the brain ^23^, and the M28L mutation likely induces a dysfunctional microglial phenotype with significant implications in AD pathology. Conversely, the P552R PLCG2 variant has been found to reduce AD risk ^9,24-27^. Therefore, determining how the M28L and P522R variants affect enzyme activity and microglial phenotype will provide valuable insights into the role of *PLCG2* in AD. Importantly, further analysis of the *PLCG2* M28L variant using human datasets is limited due to the very low frequency of this variant. Specifically, it was not possible to investigate the interaction of the *PLCG2* M28L variant with *PLCG2* expression, amyloid plaques, and *APOE* ε4. Therefore, larger cohorts are required to confirm our observation and perform a more comprehensive study of the role of the *PLCG2* M28L variant in microglia-mediated AD pathology.

Furthermore, we found that *PLCG2* expression increases in several brain regions of LOAD patients, which correlates with brain amyloid plaque density and expression levels of microglial markers *AIF1* and *TMEM119* ^*28-30*^. These results highlight an important relationship between amyloid plaques and *PLCG2* expression, which is further supported by increased levels of *Plcg2* throughout disease progression in the 5xFAD mice, a well-studied model of amyloid pathology^22^. *PLCG2* expression is also upregulated in the amyloid models TgCRND8 and App ^NL-G-F/NL-G-F^ mice^31^. Interestingly, microglia surrounding cortical plaques in TgCRND8 mice express *Plcg2* ^23^, further implicating *PLCG2* in microglial response to Aβ. Nonetheless, further studies are necessary to investigate the role of *PLCG2* in plaque-associated microglia.

eQTL analysis was performed to gain insight into the relationship between genetic variants and *PLCG2* expression levels. The SNP most significantly associated with *PLCG2* expression levels was rs4420523, an intronic variant. Within the same haplotype blocks ^32^, about ten variants that significantly increased *PLCG2* expression were strongly correlated with rs4420523, including a synonymous *PLCG2* variant, rs1143688. Further studies to investigate whether increased gene expression by the *PLCG2* variants is related to AD pathogenesis are of great interest.

In conclusion, we identified a novel association between a missense variant in *PLCG2* (M28L) and an increased risk for LOAD. Additionally, we showed that *PLCG2* expression was increased in several brain regions of LOAD patients and strongly correlated with brain amyloid burden in LOAD patients and AD animal models. Our results provide further evidence that *PLCG2* may play an important role in AD pathophysiology. Future studies investigating how the M28L and P522R variants in *PLCG2* modulate its function and microglia phenotypes in AD could lead to the identification of potential therapeutic strategies focused on PLCG2 activity.

## Materials and Methods

### Human participants and RNA-Seq

RNA-Seq and whole-genome sequencing data were obtained from the Accelerating Medicines Partnership for Alzheimer’s Disease (AMP-AD) Consortium, where all individuals used in the analysis were participants of the Mayo Clinic Brain Bank cohort, the Mount Sinai Medical Center Brain Bank (MSBB) cohort, and the Religious Orders Study and Memory and Aging Project (ROSMAP) cohort.

In the Mayo Clinic RNA-Seq dataset^33^, RNA was isolated from the temporal cortex (TCX) and cerebellum (CER). The RNA-Seq-based whole transcriptome data were generated from human samples of 151 TCX (71 cognitively normal older adult controls (CN) and 80 LOAD) and 151 CER (72 CN and 79 LOAD). LOAD had a Braak score of ≥ 4.0 and met neuropathological criteria for AD, while CN had a Braak score of ≤ 3.0 and without neurodegenerative diagnoses. The quality of the samples was selected to be RNA integrity number (RIN) ≥ 5.0 for inclusion in the study.

In the MSBB dataset^34^, RNA was isolated from the top two most vulnerable regions of LOAD, parahippocampal gyrus (PHG) and inferior frontal gyrus (IFG), and the ranked 7^th^ and 14^th^ most vulnerable brain regions of LOAD, superior temporal gyrus (STG) and frontal pole (FP).

The assessment of dementia and cognitive status was conducted using a clinical dementia rating scale (CDR)^35^. LOAD had a CDR score of ≥ 0.5, while mild cognitive impairment (MCI) had a CDR score of 0.5, and CN had a CDR score of 0 without any significant memory concern of their daily activity. This study included 108 participants (16 CN, 14 MCI, and 78 AD) for PHG, 137 participants (21 CN, 18 MCI, and 98 LOAD) for STG, 136 participants (18 CN, 16 MCI, and 102 LOAD) for IFG, and 153 participants (22 CN, 20 MCI, and 111 LOAD) for FP.

In the ROSMAP dataset^36^, RNA-Seq was performed on dorsolateral prefrontal cortex from 241 participants (86 CN and 155 LOAD). The diagnosis is based on the assessment of clinical diagnosis of cognitive status, which followed the criteria of the joint working group of the National Institute of Neurological and Communicative Disorders and Stroke and the Alzheimer’s Disease and Related Disorders Association.

### Whole Genome Sequencing (WGS)

Pre-processed whole-genome sequencing data were obtained from the Accelerating Medicines Partnership for Alzheimer’s Disease (AMP-AD) Consortium ^33,34,36-38^. Whole-genome sequencing libraries were prepared using the KAPA Hyper Library Preparation Kit per the manufacturer’s instructions. Libraries were sequenced on an Illumina HiSeq X sequencer using pair-end read chemistry and read lengths of 150bp. The paired-end 150bp reads were aligned to the NCBI reference human genome (GRCh37) using the Burrows-Wheeler Aligner (BWA- MEM)^39^ and processed using the GATK best practices workflow that marks potential duplicates, locally realigns any suspicious reads, and re-calibrates the base-calling quality scores using Genome Analysis Toolkit (GATK)^40^. The resulting BAM files were analyzed to identify variants using the HaplotypeCaller module of GATK for multi-sample variant callings^41^.

### Statistical analysis

To investigate the diagnosis group difference of *PLCG2* expression between CN, MCI, and LOAD, we used the *limma* software^42^ to perform a differential expression analysis^36^. Age, sex, and *APOE* ε4 carrier status were used as covariates. To investigate the relationship between *PLCG2* expression levels and amyloid plaque density or expression levels of microglia specific markers (*AIF1 and TMEM119)*, we used general linear models with *PLCG2* expression levels as a dependent variable and plaque density or microglia specific markers as well as age, sex, and *APOE* ε4 carrier status as explanatory variables. The regression was performed with the “glm” function from the stats package in R (version 3.6.1).

In the mouse study, statistical analyses were performed using GraphPad Prism (Version 8.4.2). Experiments at the 2-, 4-, 6- and 8-month time points were performed independently, so statistical comparisons between wild-type and 5xFAD AD mice were performed by unpaired t-test. Graphs represent the mean, and error bars denote the SEM.

### Mice

All wild-type and 5XFAD mice used in this study were maintained on the C57BL/6J background and purchased from Jackson Laboratory (JAX MMRRC Stock# 034848). All mice were bred and housed in specific-pathogen-free conditions. Both male and female mice were used in this study.

### Mouse RNA isolation for qPCR

Mice were anesthetized with Avertin and perfused with ice-cold PBS. The cortical and hippocampal regions from the hemisphere were micro-dissected and stored at -80-degree C. Frozen brain tissue was homogenized in buffer containing 20 mM Tris-HCl (pH=7.4), 250 mM sucrose, 0.5 mM EGTA, 0.5 mM EDTA, RNase-free water, and stored in an equal volume of RNA-Bee (Amsbio, CS-104B) at -80-degree C until proceeding to RNA extraction. RNA was isolated by using chloroform extraction and purified by using the Purelink RNA Mini Kit (Life Technologies). The cDNA was prepared from 750 ng of RNA by using high-capacity of RNA-to-cDNA kit (Applied Biosystems), and the qPCR was performed by using the StepOne Plus Real-Time PCR system (Life Technologies) with Taqman Gene Expression Assay (Mm01242530_m1 from the Life Technologies). The relative gene expression was determined by using ΔΔCT and assessed relative to GAPDH (Mm99999915_g1). A Student’s t test was performed for qPCR assays, comparing the results of wild-type and 5XFAD animals.

### Expression quantitative trait loci (eQTL) analysis

To perform an eQTL analysis of *PLCG2* expression levels, we used common SNPs (±20kb from the gene boundary; minor allele frequency (MAF) > 5%) from whole-genome sequencing. Only non-Hispanic Caucasian participants were selected. In addition, age and sex were used as covariates. The results of the eQTL analysis were plotted with LocusZoom ^43^.

### Homology Modeling

The model of PLCG2 structure containing all residues from amino acid 14-1190 was built by using the template of PLCG1 model ^44^. The cartoons with substitutions of M28L and P522R were generated with PyMol (The PyMOL Molecular Graphics System, Version 2.0 Schrödinger, LLC).

## Author contributions

A.P.T, J.S, S.J.B, A.L.O, G.W.C, Y.L, G.E.L, B.T.L, and K.N designed the study. A.P.T, C.D, C.P, M.M, P.B.L, N.H, and K.N performed the experiments and analyzed the data. A.P.T, M.M, G.E.L, and K.N wrote the manuscript. All authors discussed the results and commented on the manuscript.

## Acknowledgments

We thank Louise Pay for critical comments on the manuscript and Alison Goate (Icahn School of Medicine), Thomas Wingo (Emory University), and Aliza Wingo (Emory University) for discussion. This work was supported by NIA grant RF1 AG051495 (B.T.L and G.E.L), NIA grant RF1 AG050597 (G.E.L), NIA grant U54 AG054345 (B.T.L et al.), NIA grant R03 AG 063250 (K.N), and NIH grant NLM R01 LM012535 (K.N)

